# Functionality of arousal-regulating brain circuitry at rest predicts human cognitive abilities

**DOI:** 10.1101/2024.01.09.574917

**Authors:** Ella Podvalny, Ruben Sanchez-Romero, Michael W. Cole

## Abstract

Arousal state is regulated by subcortical neuromodulatory nuclei, such as locus coeruleus, which send wide-reaching projections to cortex. Whether higher-order cortical regions have the capacity to recruit neuromodulatory systems to aid cognition is unclear. Here, we hypothesized that select cortical regions activate the arousal system, which in turn modulates large-scale brain activity, creating a functional circuit predicting cognitive ability. We utilized the Human Connectome Project 7T functional magnetic resonance imaging dataset (N=149), acquired at rest with simultaneous eye tracking, along with extensive cognitive assessment for each subject. First, we discovered select frontoparietal cortical regions that drive large-scale spontaneous brain activity specifically via engaging the arousal system. Second, we show that the functionality of the arousal circuit driven by bilateral posterior cingulate cortex (associated with the default mode network) predicts subjects’ cognitive abilities. This suggests that a cortical region that is typically associated with self-referential processing supports cognition by regulating the arousal system.

## INTRODUCTION

Cognition is greatly influenced by ongoing fluctuations in arousal (Berridge and Waterhouse 2003; Aston-Jones and Cohen 2005; McCormick, Nestvogel and He 2020). Consider a student at a test – to perform well, they need to either increase their level of arousal, if they feel tired, or decrease it, if they feel stressed. In this case, the task – performing well on a test – defines the optimal level of arousal, whereas a student’s ability to adeptly shift their arousal level allows strategic control of behavioral attributes: for example, balancing the exploration-exploitation trade-off or adjusting reaction times (van Kempen *et al*. 2019; Waschke, Tune and Obleser 2019; Podvalny, King and He 2021). Accordingly, several scholars have theorized that arousal-related processes play a major role in general intelligence (Tsukahara and Engle 2021) and creativity (Suler 1980), whereas an inability to appropriately adjust arousal may severely compromise social, emotional, and cognitive aspirations (Yamamoto, Shinba and Yoshii 2014). Despite these crucial roles of top-down arousal control in human behavior, the possible arousal-regulating cortical circuits and their effect on general cognitive abilities remain poorly understood.

Brainstem locus coeruleus (LC) neurons, whose axon terminals extend throughout the entire cortex via extensive branching, regulate arousal by releasing norepinephrine (NE) and dopamine (DA) (Poe *et al*. 2020; Ranjbar-Slamloo and Fazlali 2020; Breton-Provencher, Drummond and Sur 2021). While evidence for LC receiving inputs back from cortex in humans is still inconclusive (Szabadi 2013), previous research points to the importance of investigating such a possibility. Electrical stimulation of dorsomedial prefrontal cortex (dmPFC) in rats, for example, results in increased LC firing, whereas chemical inactivation results in inhibition (Jodo and Aston-Jones 1997; Jodo, Chiang and Aston-Jones 1998). While LC plays a major role in arousal state regulation, other neuromodulatory systems, such as cholinergic, dopaminergic, and serotonergic systems, are also involved. In humans, Granger causality analysis of fMRI signals suggests a directional effect of posterior inferior parietal lobule (pIPL), a DMN (default mode network) region, on the arousal state (Yellin, Berkovich-Ohana and Malach 2015). These findings point to a strong possibility of cortical neurons projecting to arousal-regulating nuclei and potentially controlling their functionality.

While fMRI is often considered too sluggish to identify directional relations, tonic (resting state) neuromodulatory effects of noradrenergic projections are known to occur at a timescale slow enough to be detected and distinguished from fast glutamatergic effects of cortical projections. First, NE receptors are metabotropic – that is, requiring multiple metabolic steps for activation, which may take hundreds of milliseconds to minutes. This may explain the findings that extracellular NE in cortex, due to either tonic LC firing or direct application, results in only sluggish (seconds to minutes) effect on spontaneous brain activity in animals (Waterhouse and Navarra 2019), and the recovery from the effects of NE application in rat cortex takes up to a minute (Armstrong-James and Fox 1983). Second, noradrenergic axons lack myelin and exhibit variable conduction velocity (Aston-Jones, Segal and Bloom 1980; Berridge and Waterhouse 2003), thus target cortical cells are triggered with low temporal precision by slow tonic LC activity at resting state (1-6Hz) (Aston-Jones and Bloom 1981; Vazey, Moorman and Aston-Jones 2018). These slow timescales are very likely within the detection range of fMRI with a temporal resolution of 1 Hz.

In addition, the distinct neurobiology of ascending and descending projections to LC predicts a likely opposing functional connectivity during resting state. First, NE is considered a modulator of a neural gain – that is, it may contribute to both the suppression of spontaneous baseline activity and to the increase of relative stimulus-triggered activity (Servan-Schreiber, Printz and Cohen 1990; Aston-Jones and Cohen 2005). This is consistent with previous studies that show suppressive effects of pupil-linked arousal in human cortex (Yellin, Berkovich-Ohana and Malach 2015), and a decrease in spontaneous firing due to application of NE in rat cortex (Armstrong-James and Fox 1983; Bassant, Ennouri and Lamour 1990; Waterhouse and Navarra 2019). Further, because cortical long-range projections are mostly excitatory, a positive effect of cortical firing on LC is expected (Figure 1). Crucially, because it is unclear to what extent animal models have sufficient top-down control of the arousal state, using fMRI in healthy human volunteers to test the hypothesis of bi-directional connectivity between cortical activity and the arousal state is our best option at this time.

**Figure 1.**
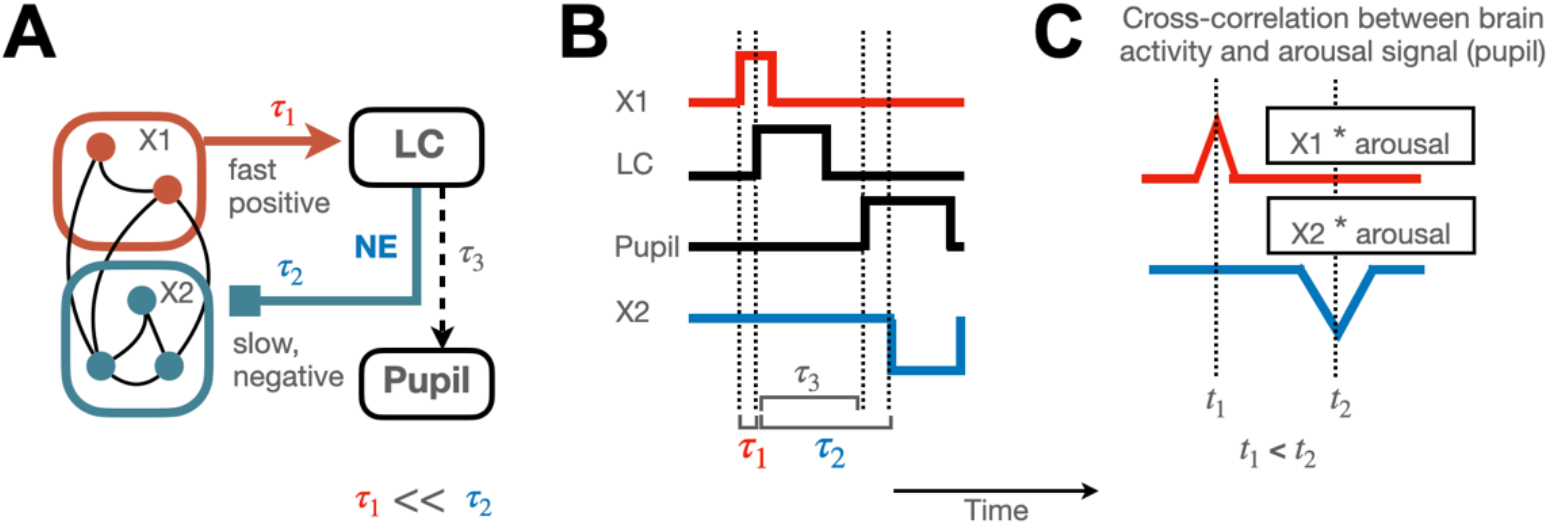
Hypothesized large-scale circuit regulating tonic arousal, based on temporal lag and sign of time series cross-correlations. **A**. Simplified wiring diagram of a large-scale brain circuit for recruitment of the brainstem arousal system. Select cortical regions (X1) signal LC via fast myelinated projections (with τ_1_ signifying temporal delay), whereas LC activation leads to slow NE release in cortex, which in turn modulates cortical sustained activity (with a delay of τ_2_). In resting state, NE release suppresses spontaneous brain activity. Non-light-mediated pupil size fluctuations reflect LC activity at rest (with a delay of τ_3_). **B**. Neural activity cascade expected from circuit depicted in A during resting state. Cortical regions X1 cause increase in LC tonic activity. X1 regions may also cause network-level activation in regions X2 (not depicted). LC tonic firing in turn results in large-scale slow release of NE, resulting in a decrease of cortical spontaneous activity with a delay of multiple seconds. **C**. Expected cross-correlation (a measure of signal similarity as a function of temporal lag) between X1/X2 type of brain regions and the arousal signal, whereas *t*_1_ and *t*_2_ depict the relative timing of the cross-correlation peaks/troughs. A positive peak is expected to appear earlier than negative trough (*t*_1_ < *t*_2_), irrespective of whether the pupil size or LC activity is used to indicate the arousal state.

Here, we aim to: (1) identify the cortical regions that exert top-down regulation of the arousal state (pupil size) and cause a subsequent large-scale activity modulation, (2) test whether the arousal-related circuit-level connectivity of each candidate region predicts individual differences in cognitive performance. To achieve these aims, we use a 7T fMRI dataset provided by the Human Connectome Project, which includes prolonged resting state fMRI recordings in a substantial number of subjects who also underwent extensive cognitive testing, allowing us to test the relationship between individual brain connectivity measures and behavioral performance.

## METHODS

### fMRI data selection and preprocessing

We used the Human Connectome Project (HCP) 7 Tesla dataset acquired at the Center for Magnetic Resonance Research at the University of Minnesota. The detailed description of the dataset is widely available – here, we provide only succinct summary pertaining our study for the readers’ convenience. 184 subjects were scanned, however, resting state data with concurrent eye-tracking were missing in 8 subjects. All subjects were generally healthy young adults between 22 and 36 years old (mean age = 29.4, standard deviation = 3.3). There were 106 females and 70 males. Resting state runs of 15 minutes (up to four per subject) were collected using gradient-echo-planar imaging (EPI) sequence with the following parameters: repetition time (TR) = 1000 ms, echo time (TE) = 22.2 ms, flip angle = 45 deg, field of view (FOV) = 208 × 208 mm, matrix = 130 × 130, spatial resolution = 1.6 mm3, number of slices = 85, multiband factor = 5, image acceleration factor (iPAT) = 2, partial Fourier sampling = 7/8, echo spacing = 0.64 ms, bandwidth = 1924 Hz/Px. The direction of phase encoding alternated between posterior-to-anterior (PA) and anterior-to-posterior (AP). During rest runs, subjects were instructed to keep their eyes open and maintain relaxed fixation on a projected bright crosshair on a dark background.

Out of 176 subjects with available data, we selected 296 sessions from 108 subjects that had high-quality eye tracking (see below selection criteria). We used “fix-denoised” data preprocessed by the HCP team (files named *_Atlas_MSMAll_hp2000_clean.dtseries.nii). Briefly, the preprocessing pipeline followed standard preprocessing procedure (motion correction, distortion correction, high-pass filtering, and nonlinear alignment to MNI template space (Glasser *et al*. 2013) plus regression of 24 framewise motion estimates (six rigid-body motion parameters and their derivatives and the squares of those) and regression of confound timeseries identified via independent components analysis (Salimi-Khorshidi *et al*. 2014; Griffanti *et al*. 2017). The data were aligned using multimodal surface matching algorithm II (MSMAII), which provides markedly better cross-subject cortical alignment than does volume-based alignment (Coalson, Van Essen and Glasser 2018).

### Eye-tracking data selection and preprocessing

Pupil diameter was utilized to estimate the within-session arousal state fluctuations. Resting state eye-tracking data were downloaded from the following HCP database directory: subj#/unprocessed/7T/rfMRI_REST*/LINKED_DATA/EYETRACKER/. Eye-tracking data in these directories were found only for 149 out of 184 subjects. In total eye-tracking data from 576 runs were downloaded (15 minutes each), 1-4 runs per subject. The eye-tracking data were acquired at 500Hz sampling rate in 68 runs and 1000Hz in 508 runs. The recordings were monocular, where left-eye pupil size data were present in 561 runs, and right-eye data in 4 runs, whereas pupil size data were completely missing from 11 runs. Considering normal blink rate of up to 15 blinks per minute, and blink duration up to 0.5 s, we normally expect up to 12.5 % (0.5 s * 15 / 60 s) pupil size data missing from the recordings. In addition, pupil size data can be missing because of uninstructed eye closure, saccades, failure of pupil detection in eye-tracking software. We allowed additional 7.5% missing data from such causes and selected only runs with pupil size data available at more than 80% of time samples, resulting in 334 runs, (supplementary figure 1A-B for missing data % distribution across runs, and examples of raw data quality) for further analysis. In addition, runs with incomplete fMRI data acquired (TR count less than 900, according to the provided session summary csv files) were excluded. Thus, in total, 296 runs from 108 subjects were selected for further analysis.

For each individual session, missing pupil data periods along with +/-100 ms surrounding samples were interpolated using Piecewise Cubic Hermite Interpolating Polynomial (PCHIP), which uses monotonic cubic splines to find the value of new points. Next, pupil size samples matching each MR image acquisition window were averaged, the timecourse convolved with a hemodynamic response function (HRF) (supplementary figure 1C-D), and z-scored to create the pupil-linked arousal predictor (depicted in Supplemental Figure 1).

### Cross-correlation analysis and peak/trough estimation

We conducted a cross-correlation analysis, which quantifies the level of synchrony between the pupil size predictor (see above and Supplemental Figure 1) and fMRI activity at consecutive temporal delays as follow:

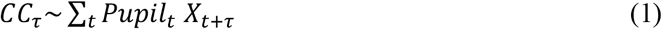

Where CC stands for cross-correlation, Pupil for pupil-linked arousal predictor, X indicates resting-state fMRI activity at a brain region, τ indicates the time delay and t indicates the collected time sample. Each point of the cross-correlation function is calculated via introducing a time delay between the two timecourses (pupil and brain) and calculating their product. A high cross-correlation value at a negative time lag is consistent with brain activity causing a change in pupil-linked arousal. Note, however, since the delay between locus coeruleus and pupillary response at resting state cannot be calculated precisely (see Figure 1A), it is difficult to interpret the values that are close to a time lag of zero. We tested the group-level cross-correlation effect via a one-sample t-test with a null distribution created by reversing the timecourse of the pupil size. Finally, for each brain region we detected a significant minimum (trough) and maximum (peak) point which was used in the following analyses.

### Receptor density and arousal-linked spontaneous activity inhibition

We used receptor density maps calculated from microarray data provided in the Allen Human Brain Atlas (Gryglewski *et al*. 2018; van den Brink, Pfeffer and Donner 2019). The maps of neuromodulatory receptors were parcellated according to the Glasser atlas (Glasser *et al*. 2016) and used as predictors of arousal-linked inhibition of spontaneous brain activity as follow:

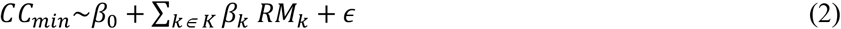

Where *CC*_*min*_ indicates the subject average minimal cross-correlation between pupil size and brain activity, and *RM* indicates the Receptor Map of receptor density data. Only regions showing significant cross-correlation values were used, which included 346 out of 360 Glasser brain regions. K=48 receptor maps were used (see supplementary figure 3), including multiple variants of noradrenaline (norepinephrine), acetylcholine, dopamine, serotonin and histamine receptors.

### Identification of arousal driver regions

We used a two-step procedure to ensure acceptable computational complexity required for model convergence. First, we constructed a network-specific linear mixed model for arousal predictor *Pupil*, as follows:

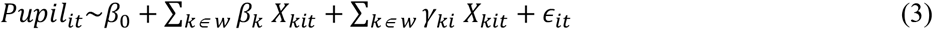

Where *X* denotes the timecourse of brain activity in potential driver region *k* within one resting state network *w, β* represent fixed effects and *γ* random effects, *i* is the subject and *t* is time in TR units. The analysis was conducted separately for each hemisphere and included *n* = 108 participants, whereas each participant’s dataset contained between 900 to 3600 time points, resulting in 265,500 total observations for each model variable. No network-specific model contained brain regions with variance inflation factor larger than 10, indicating acceptable multicollinearity. Only random slopes were included since brain activity was normalized for each participant to ensure standardized parameters estimates. To allow reliable model fit and ensure convergence, the timing of maximal cross-correlation between individual brain region’s activity and pupil size was calculated, and the brain activity shifted in time assuring optimal alignment. Finally, as the second step of the procedure, the regions that showed significant fixed effects for each network were then submitted to a cross-network linear mixed model (same formula as above).

### Mediation analysis

Mediation analysis was conducted to test whether driver regions’ effects on cortical network activity is partially explained via pupil-linked arousal. For each driver region and each network or subcortical region, we conducted a group-level mediation analysis using two linear mixed models:

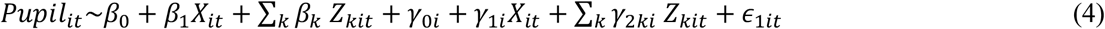

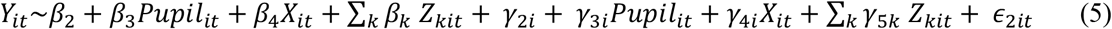

Where *X* represents the driver region (identified according to eq. (3)) activity and *Y* represents the outcome network averaged activity for subject *i* at time *t. Z* represents activity of remaining driver regions within the same hemisphere. To calculate *Y*, the region-level brain activity was first shifted according to the time of strongest negative correlation with pupil size, and averaged across all regions. (As a result of this procedure, the trough of the correlation between the averaged brain activity and pupil size was centered at zero.) *βs* are fixed effects parameters (shared by all subjects), *γ*_*i*_ are random effects parameters for subject *i*. The mediation effect was calculated as ACME (Average Causal Mediation Effect), which equals the product of *β*_1_ and *β*_3_ The *p*-value of ACME was estimated via bootstrapping of 1000 simulations using the Statsmodels toolbox, which uses independent random sampling of exogenous and endogenous variables. A significant mediation effect (ACME) indicates that the effect of a driver region on network activity can be partially explained by pupil-linked arousal.

### Cognitive ability scores

We used fluid cognition composite scores calculated from NIH toolbox Cognition Battery in conjunction with Penn matrices reasoning test scores to assess individual cognitive ability. The fluid cognition composite score is computed as an aggregate of behavioral metrics in the Dimensional Change Card Sort (DCCS) test, Flanker, Picture Sequence Memory, List Sorting, and Pattern Comparison measures. Briefly, in the DCCS test the participant is asked to match a series of picture pairs to a target picture according to stimulus dimensions that change across trials (color or shape) and is used to assess cognitive flexibility. In the Flanker test the participant is asked to focus on a particular stimulus while inhibiting attention to the stimuli flanking it—this test is used to assess inhibitory control and attention. The Picture Sequence Memory test assesses episodic memory—participants are shown pictures of objects and activities, and then asked to reproduce the sequence of pictures as it was presented to them. List Sorting is a working memory test, which requires participants to recall and sequence different visual and auditory stimuli. The Pattern Comparison test specifically targets the measurement of processing speed—the participants are requested to determine whether two stimuli are the same or not. To calculate the aggregate, the test results were first converted to normally distributed scores and then averaged across tests. More details on measures and their validity are available in (Heaton *et al*. 2014; Scott, Sorrell and Benitez 2019). We used age-unadjusted scores and opted for including age as a variable in the model because of the previously reported interaction between arousal dynamics and age (Liu *et al*. 2020). Fluid cognition composite scores were averaged with a performance metric (number of correct responses) from the Penn matrices reasoning test (both metrics z-scored prior to averaging). This test was originally designed as a simplified Raven’s Standard Progressive Matrices test including only 24 items (unlike 60 in Raven’s) and includes abstract geometric patterns to be completed via multiple choice options.

### Analysis of individual differences in behavior

To examine individual differences in behavior, we first fit a mediation model for each subject as follow (analogously to a group-level model):

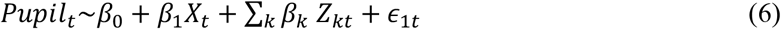

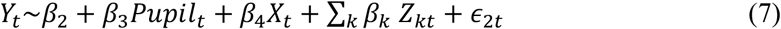

Next, we use the product of *β*_1_ and *β*_3_ (that is, ACME) averaged across networks or subcortical regions to predict the behavioral variables. Because both fluid reasoning ability and arousal dynamics are affected by age, we also included age as a covariate in the analysis:

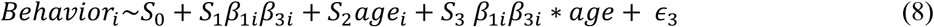

Control models, included *β*_1_or *β*_3_, and age:

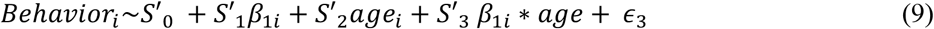

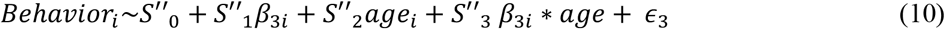

## RESULTS

Our first goal was to identify cortical regions that are likely to regulate the arousal system (Fig.1). We reasoned that activity in such regions must satisfy (at least) the following conditions:

i. <u>Directional effect</u>: Brain activity in cortical arousal-driver regions must be positively correlated with an indicator of arousal (pupil size) since the long-range projections from cortex to LC are more likely to be excitatory (see *Introduction* above for details) (Fig. 1);
ii. <u>Independent effect</u>: it is possible that two cortical regions, A and B, predict the arousal signal, while in fact only region A is connected to LC whereas region B is merely connected to A. For this reason, to determine the effects of cortical activity on the arousal signal, we need to account for correlations among cortical regions.
iii. <u>Circuit-level effect:</u> It must be demonstrated that the arousal driver regions induce delayed negative effects on spontaneous brain activity (due to NE-linked suppression) that are specifically *mediated* by the arousal signal (i.e., cannot be fully explained by direct cortical-cortical connections).

While it is theoretically possible that the above procedure will misidentify cortical regions that are affected by activity from regions we do not measure (“unmeasured confounders2), this is unlikely with the full brain coverage of fMRI. It is also likely that additional cortical regions that suppress the arousal system via the generally rare long-range inhibitory projections or by targeting inhibitory interneurons within LC exist in the brain. However, we do not test for such a hypothesis herein since such anatomically fine-grained effects are difficult to discern with fMRI, where each voxel contains hundreds of thousands of neurons.

### Select cortical regions exhibit the properties of arousal regulation

To identify the potential arousal driving regions we used 7T fMRI HCP data recorded at 1 Hz temporal resolution during resting state with concurrent eye tracking (see Methods). We opted to using the pupil size as the arousal signal rather than LC activity since the signal acquisition methodology was not specifically optimized for LC in the HCP dataset, and non-light-mediated ongoing fluctuations of pupil size have been shown to reflect resting state electrophysiological LC activity with temporal precision higher than that of hemodynamic response.

We first examine the cross-correlation between fMRI BOLD signal and pupil size for each brain region (Figure 2A-B). The results include positive correlations between multiple brain regions and pupil size, at a negative time lag (i.e., brain activity precedes the change in pupil size), and thus satisfying the first criterion for arousal-driving regions (“directional effect2). Note that the regions positively correlated with pupil size are situated mostly in frontal and parietal brain regions belonging to default mode network (DMN), frontoparietal network (FPN) and cingulo-opercular network (CON) (Figure 2A-B, supplemental figure 2 or figure 2E for network partition). The positive correlation peak appears significantly earlier than the correlation trough (Figure 2B, 2D), which dominates the brain networks specializing in motor responses and processing of sensory stimuli (e.g., Vis2, SMN). We confirm that such negative correlations are consistent with NE-driven inhibition of spontaneous activity via testing the predictive power of neuromodulatory receptor distribution on the arousal-linked inhibition effect (see Supplementary figure 3 and Methods). While the density of ADRA1B (NE) receptor is the strongest predictor of arousal-linked inhibition, other receptors, such as cholinergic receptors, seem to also play a role, whereas dopamine D4 receptor density predicts the opposing effect (less arousal-linked inhibition with higher density). Further, this result is consistent with the possibility of suppression of spontaneous brain activity due to increase in neural gain (Figure 2C) and with previously reported correlations with pupil size that were positive in DMN and negative in visual brain areas (Yellin, Berkovich-Ohana and Malach 2015).

**Figure 2.**
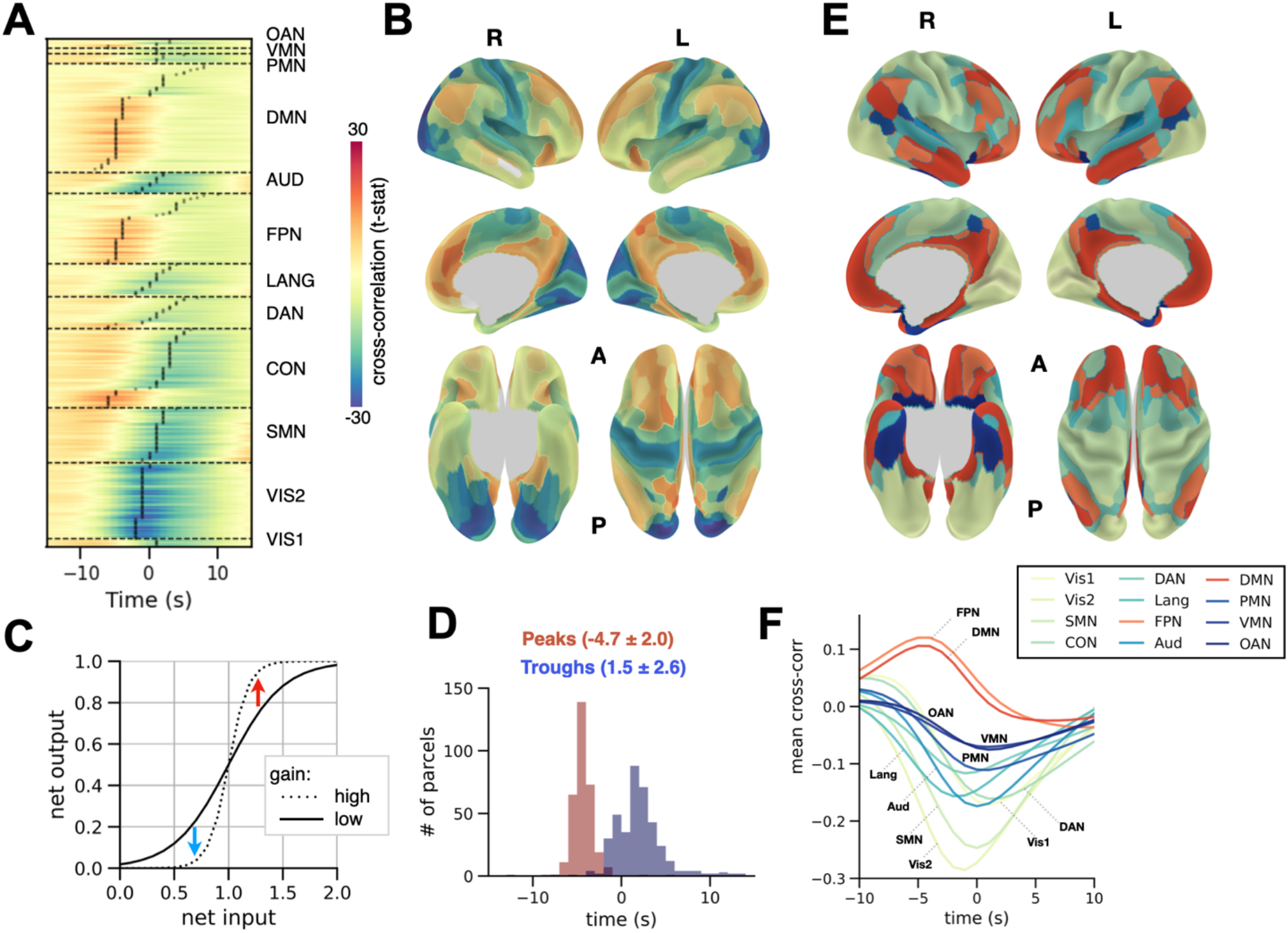
Spatiotemporal interactive dynamics of arousal state and spontaneous brain activity. **A**. Cross-correlation between pupil size predictor and fMRI cortical parcel-level activity (Glasser parcellation) compared with a null distribution calculated via reversal of pupil size time course (t-statistic). The horizontal axis depicts the temporal delay at which the correlation was calculated. The parcels are grouped according to large-scale resting-state brain networks (Ji *et al*. 2018) and sorted according to peak/trough time delay (black dots). **B**. Peaks/trough amplitude of pupil-brain cross-correlation (same color scale as A) presented on the cortical surface. **C**. Illustration of gain modulation depicting the potential suppression of low brain activity such as during resting state (blue arrow) and amplification of high brain activity (red arrow). **D**. Distribution of cross-correlation peak/trough time lags across brain regions. **E**. Large-scale resting-state networks presented on the cortical surface. **F**. Averaged cross-correlation between pupil size and brain activity within each resting-state network (Vis1:Visual 1, Vis2: Visual 2, SMN:somatomotor, CON:cingulo-opercular, DAN:dorsal attention, Lang:language, FPN: frontoparietal, Aud:Auditory, DMN: default mode, PMN:posterior multimodal, VMN: ventral multimodal, OAN: orbito-affective). Networks that on average show larger absolute value of positive peak than negative trough are colored in warm colors (FPN and DMN), same color code as (**E**).

The positive cross-correlation between region-level brain activity and pupil size could either reflect arousal-driving cortical regions or merely reflect propagation of activity within cortical resting state networks without causally affecting the arousal state (see criterion (ii)). To discern between these two possibilities, we constructed a two-step model that progressively eliminated cortical regions that lost the effect on arousal due to cross-region correlation (see Methods). In the first step, we construct a model for each resting state network to test the effect of each candidate brain region on the arousal state while including all other regions within the same network as covariates. In the second step, we combine all regions that show significant effects in the multiple models of the first step to account for possible cross-networks correlations (see Methods, Equation 1). As expected, this method eliminated most cortical brain regions that showed a positive effect on the arousal state regardless of network-level correlations (compliant with criterion (i), Figure 2B) and only the regions that showed significant *independent* effects on arousal state remained (compliant with criterion (ii), presented in Figure 3A).

**Figure 3.**
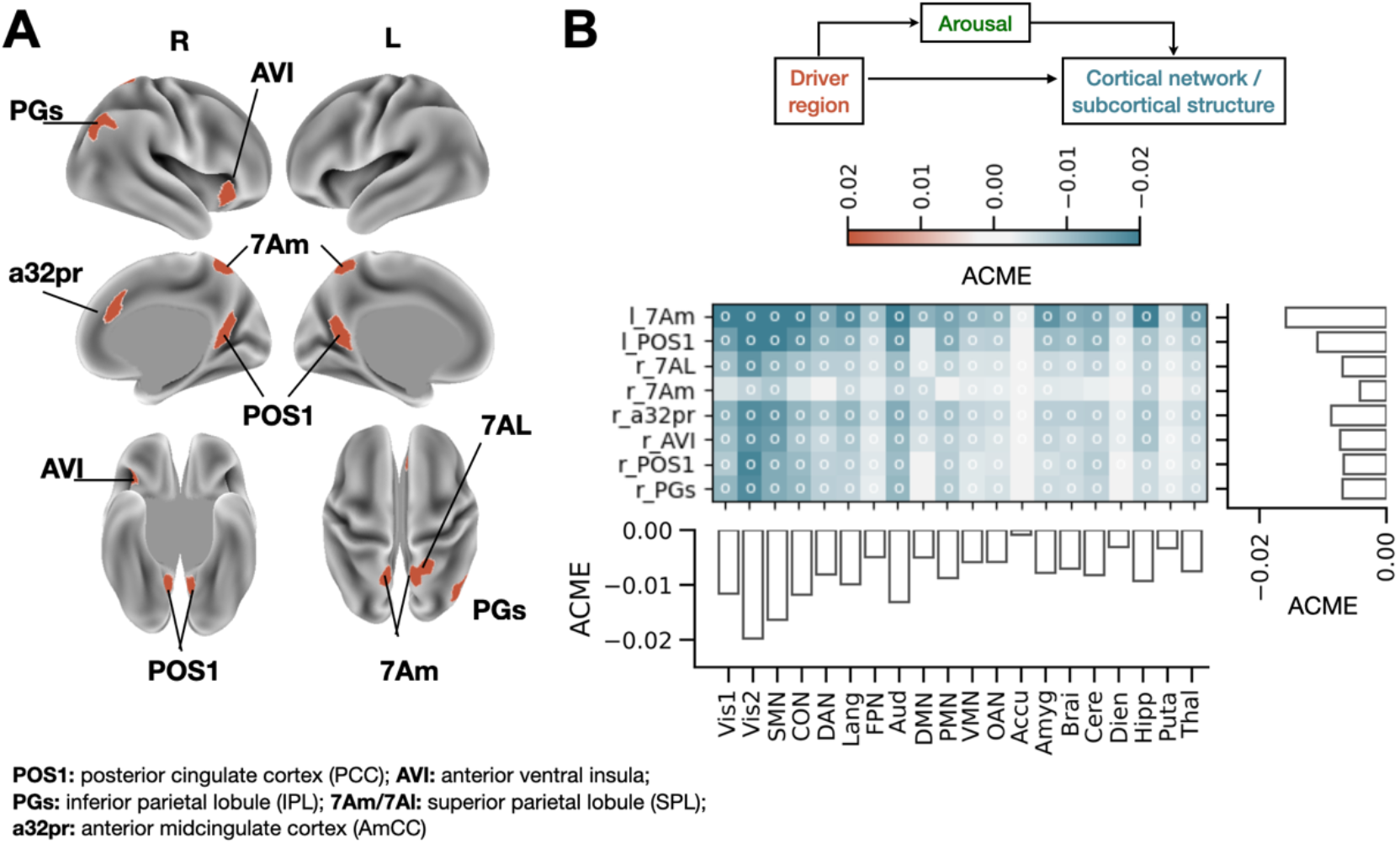
Regions identified as regulating large-scale spontaneous activity via pupil-linked arousal. **A**. Brain regions (Glasser parcellation) predictive of pupil-linked arousal in agreement with “independent effect2 criterion. These regions were identified by considering network-level and cross-network activity correlations via multiple regression linear mixed models. The legend provides commonly used names for Glasser brain regions. **B**. Effects of brain regions presented in A on large-scale network and subcortical activity as mediated by arousal. **Top:** illustration of causal mediation analysis conducted for each region presented in A. **Bottom:** Each cell of the matrix shows ACME (Average Causal Mediation Effect), representing the level of influence of brain activity in the arousal-driver region on subsequent activity in the cortical network or subcortical structure that is specifically routed via arousal. The white circles indicate p-value < 0.05 (FDR corrected). The bar plots show the ACME averaged across the arousal-driver regions (horizontal bars) and across the arousal-driven networks/subcortical areas (vertical bars). The network abbreviation is the same as Figure 2.

The candidate arousal-driver regions identified (Figure 3A) include: (1) bilateral POS1 (posterior-occipital sulcus 1), regions residing within the posterior cingulate cortex (PCC); (2) bilateral 7Am (7 anterior-medial), regions in superior parietal lobule located on the medial side; (3) right 7AL (7 anterior-lateral), found also in superior parietal lobule on the lateral side; (4) PGs (parietal area G superior) – area in the angular gyrus of inferior parietal lobule (IPL); (5) a32pr (anterior 32 prime), a region in anterior midcingulate cortex; and (6) AVI, anterior ventral insular area.

To test whether these candidate arousal-driver regions (Figure 3A) indeed invoke large-scale activity modulation via the arousal system (criterion (iii)), we conducted a causal mediation analysis. Namely, we used linear mixed effect models testing the effect of each driver region on brain-wide activity via arousal state (pupil size) while the potential direct effect (bypassing arousal) is accounted for (Figure 3B top, see Methods, Equations 4-5) (Figure 3B). According to our hypothesis, we expect to see a negative mediation effect because the arousal driver regions should induce inhibition of spontaneous brain activity via the arousal system. This analysis revealed that all candidate arousal-driver regions showed significant mediation effects (figure 3B bottom, FDR corrected for multiple comparisons). Note the regions of left superior parietal lobule (left 7Am) and PCC (POS1) showed the strongest effects, suggesting these regions may play an especially prominent role in regulating widespread cortical activity via the arousal system. Direct effects (driver-to-network) were not eliminated by including the pupil size as a mediator, hence the mediation effects reported herein can only be considered partial (as expected due to extensive cortical-cortical connectivity).

### Large-scale arousal circuit connectivity predicts individual cognitive performance

In a final analysis, we tested the hypothesis that the identified large-scale arousal-regulating circuits predict individual differences in cognitive performance. This hypothesis stems from the idea that each cognitive task requires an arousal level that is optimal for that specific task; thus, an adaptive arousal-regulating circuit should improve general cognitive ability rather than one specific task. To this end, we calculated the cognitive ability score for each participant as an aggregate performance metric across cognitive tasks (see Methods, Figure 4A). Next, we fit a mediation model for each individual subject (see Methods) to estimate the subject-level average causal mediation effect (ACME) for each arousal-driving and arousal-affected brain area. The distribution of ACME averaged across brain regions is depicted in Figure 4B. The negative values of ACME signify arousal-driven suppression of spontaneous brain activity. The relationship between global ACME and individual cognitive ability was negative (coef. = - 27.5, p-value = 0.043, Figure 4D).

**Figure 4.**
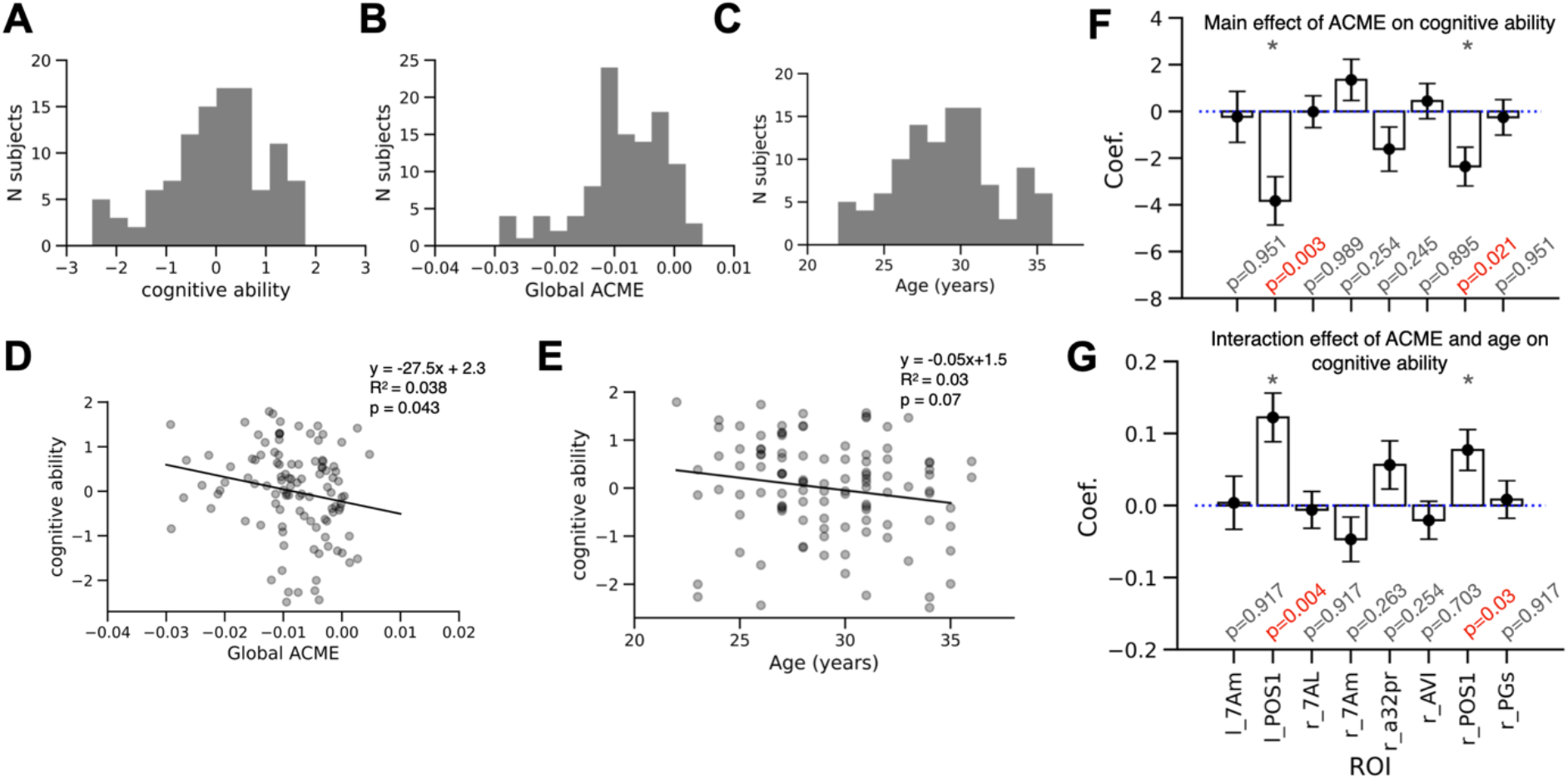
Functionality of arousal-regulating brain circuit predicts individual differences in cognitive ability. **A**. Distribution of cognitive ability scores across subjects. **B**. Distribution of global ACME (Averaged Causal Mediation Effect), that is, ACME averaged across the driving and the affected brain areas. **C**. Distribution of age across subjects. **D**. The relationship between global ACME and cognitive ability with linear model fit result depicted and inferred parameters specified. **E**. The relationship between participant age and cognitive ability. **F**. Main effect of ACME, signifying the functionality of the large-scale arousal circuit driven by each candidate arousal-driving region on cognitive ability. P-values are specified after FDR correction and the values below 0.05 are marked in red. **G**. Same as F but for interaction effect of ACME and age on cognitive ability. The main effect of age was not significant for any of these regions.

Previous research has shown that both LC integrity (Liu *et al*. 2020) and cognitive performance (Bugg *et al*. 2006) decline with age. In our data, the participants’ age was between 22 and 36 (Figure 4C), and the effect of age on cognitive ability, while not statistically significant (coef. = -0.05, p-value = 0.07, Figure 4E), is consistent with the previous findings.

Because we identified brain regions that independently drive the arousal state, we are especially interested in testing which region-specific arousal-related circuits predict cognitive ability. To this end, since arousal-driven suppression of spontaneous brain activity is global, for each arousal driver region we averaged the causal mediation effect across the brain (Figure 3B, right horizontal bars). Next, we tested whether this effect (ACME), signifying arousal-circuit functionality, predicted cognitive ability. We also included age as a covariate and as an interaction term with ACME in our next model (Methods, Equation 8).

Individual-subject scores of cognitive ability were significantly predicted by the circuit-level connectivity of left (coef. = -3.8, p-value = 0.003 [FDR-corrected], BIC [Bayesian Information Criterion] = 307.9) and right (coef. = -2.4, p-value = 0.021, BIC = 312.9) POS1 (PCC) only (Figure 4F). The effect was negative, indicating that stronger arousal-linked control by PCC predicts a higher cognitive ability score (since the mediation effect gets stronger in the negative direction). The effect of age on cognitive ability scores was negative, which is consistent with prior literature, but again not significant (p-value > 0.05) in the relatively narrow age group that was included in the present study. We also detected a positive interaction effect with age in the bilateral POS1 (coef = 0.12, p-value =0.004 and coef = 0.08, p-value =0.03) (Figure 4G), indicating that the effect of the arousal circuit on cognitive performance gets weaker with age.

Finally, as an exploratory control analysis, we tested whether the effects above could be explained via individual pathways from driver regions to arousal or from arousal to the rest of the brain (see Methods, Equations 9 and 10 correspondingly). The model testing the pathway of the driver region on arousal state without considering arousal-linked inhibition of network activity (eq. 9) showed significant effects in both left and right PCC (coef = 2.21, p-val = 0.03, BIC = 313.9 and coef = 2.37, p-val = 0.02, BIC = 312.2). By contrast, the model testing the pathway from arousal state to global inhibition of spontaneous brain activity (eq. 10) did not produce significant results (p > 0.05). As an additional exploratory analysis we tested whether an effect of arousal-linked spontaneous activity inhibition in any particular network had a predictive effect on behavior – the results were not significant after FDR correction. However, the model of arousal-linked inhibition in Vis2 network showed the best predictive power (coef. = -2.00, BIC = 313.4) with a p-val = 0.009 before FDR-correction; Note that Vis2 network also shows the strongest arousal-linked inhibition on average (see figure 2A-B)). Together, these results confirm that the model using ACME of the right PCC and global arousal inhibition (Figure 4F) provides the lowest Bayesian Information Criterion, which suggests this model should be preferred for prediction of cognitive performance.

## DISCUSSION

Prior research regarding the arousal system has mainly focused on the investigation of how the ascending pathway of this system shapes cognitive task performance. By contrast, the questions of whether and how higher-order cortical regions regulate the arousal state in a top-down manner has remained underexplored. Here we hypothesized that the quality of such top-down regulation in individual subjects is predictive of their cognitive abilities because each cognitive operation benefits from distinct level of arousal. The present work sheds light on the mechanisms underlying the cortical regulation of the tonic arousal state (inferred via pupil size): First, we identify select cortical regions that independently predict the arousal state (Figure 3A) – specifically, these regions are situated in posterior cingulate cortex (PCC), anterior ventral insula (AVI), superior parietal lobule (SPL), inferior parietal lobule (IPL), and anterior midcingulate cortex (AmCC); Second, we show that activity in these regions predicts large-scale cortical state modulation that is specifically mediated by the arousal system (Figure 3B); We confirm that arousal-linked inhibition is predicted by density of neuromodulatory receptors (Supplementary Figure 3). Finally, we show that the properties of one such identified arousal-regulating neural circuits – involving the PCC – predicts individual differences in an aggregate cognitive performance metric (Figure 4), in agreement with our hypothesis.

### Control of the arousal system by frontal and parietal cortical regions

PCC and IPL are interconnected regions within the default mode network (DMN) (Buckner, Andrews-Hanna and Schacter 2008). While DMN activity has been traditionally considered task-irrelevant, recent research provides evidence that PCC is active in vast variety of tasks (Pearson *et al*. 2011) (Foster *et al*. 2023) – for example, visuospatial orientation and planning (Vatansever *et al*. 2018), orientation in time and inference of social relationships (Peer *et al*. 2015), subjective value representation (Kable and Glimcher 2007; Levy *et al*. 2010), and emotional processing (Maddock, Garrett and Buonocore 2003). Further, anticipatory prestimulus activity in PCC mediates allocation of top-down spatial attention (Small *et al*. 2003) and predicts subsequent insight during a problem solving task (Kounios *et al*. 2006). Previous work also suggests that a small number of DMN regions, and PCC in particular, are recruited when a change of context is required (Crittenden, Mitchell and Duncan 2015; Smith, Mitchell and Duncan 2018). According to our results, one plausible explanation of DMN involvement in such an overwhelming variety of tasks is that DMN regions have the capacity to regulate tonic arousal.

Our data suggests that bilateral PCC (POS1) and right IPL (PGs), despite being interconnected anatomically and functionally, show independent effects on the arousal state at rest (Figure 3A). This result is consistent with a previous human 3T fMRI study examining the interaction between cortical activity and the arousal state in 20 individuals (Yellin, Berkovich-Ohana and Malach 2015). In our study of 107 individuals with 7T fMRI, however, ventromedial prefrontal cortex (vmPFC), another major DMN hub, despite showing a positive correlation with arousal (Figure 2B), did not show a significant group-level independent effect (Figure 3A). Interestingly, we find that a region in anterior midcingulate cortex, which is situated in close proximity to vmPFC, does show an independent contribution to predicting the tonic arousal state. Such a discrepancy could be explained by lack of control for within-network functional connectivity in the previous study, although smaller sample size, lack of cortical parcellation and group-level differences between subjects recruited in the two studies are also likely to affect the results.

Sleep studies in humans show that PCC and IPL activity is positively correlated with wakefulness (highest during wakefulness and lowest during sleep) (Maquet 2000; Vogt and Laureys 2005; Sämann *et al*. 2011), similarly to LC tonic firing rate in animals, which is also highest in states of stress and lowest in sleep (Aston-Jones and Bloom 1981; Rasmussen, Morilak and Jacobs 1986; Berridge and Waterhouse 2003; Takahashi *et al*. 2010). Interestingly, the correlation between PCC and mPFC that is normally observed in resting state disappears in deep stages of sleep (Horovitz *et al*. 2009). These prior findings are consistent with the lack of independent contribution of mPFC to predicting the arousal state that we observe in our resting state data.

In addition to the DMN regions we discuss above, we have also identified five additional regions (Figure 3A) in frontal and parietal cortices that show properties of arousal regulation. While these five regions are not part of the DMN, they are situated in close proximity to the DMN. The properties of arousal-regulating circuits involving these regions did not predict individual differences in cognitive performance (Figure 4F), yet they may play role in behaviors that were not tested in the present study.

Right anterior ventral insula (rAVI), which is a part of FPN, has been mostly implicated in processing of affective experience (Wager *et al*. 2008; Touroutoglou *et al*. 2012; Uddin 2015) and is suggested to play a role in present-moment awareness and control of autonomic nervous system processes(Craig 2009). Our results are consistent with the idea that the rAVI recruits the brainstem arousal system to regulate tonic arousal state underlying the processing of emotions. One should note, however, that this is a cytoarchitectonicaly complex area only recently parcellated from anterior insula (Baker *et al*. 2018a), which does not have a topological equivalent in macaques, mice or rats(Uddin 2015; Namkung, Kim and Sawa 2017). For these reasons, the previous studies may need be reevaluated and investigation of this area in humans is especially crucial. For example, the adjacent dorsal anterior insula area, belonging to CON, has been previously proposed to play a role in tonic alertness regulation during task performance (Sadaghiani *et al*. 2015) and therefore dedicated future experiments are needed to disentangle between the distinct components of emotion-related and emotion-neutral tonic arousal state regulation, along with recordings from both ventral and dorsal insula and LC.

Anterior midcingulate cortex (AmCC, a32pr) has been previously implicated in evaluating motivation, anticipating outcomes, recognizing reward values(Bush, Luu and Posner 2000; Bush *et al*. 2002) as well as proactive prediction and monitoring of decision outcomes during social interactions(Apps, Lockwood and Balsters 2013). Such involvement in proactive and lasting behavioral modulation is consistent with our findings of this area affecting tonic arousal. Further, while AmCC has no cytoarchitectonic equivalent in rats, mice, and macaques (Vogt 2016), rodent ACC is considered to be a close relative and anterograde tracing of rat ACC axons shows their close proximity to distal LC dendrites (Gompf *et al*. 2010), which is also consistent with our findings.

Superior parietal lobule (SPL, 7am/7al) is an area that is involved in multimodal integrated visuo-spatial representation, egocentric body coding and attentional processes (Scheperjans *et al*. 2008; Wang *et al*. 2015). 7al belongs to the somatomotor network and shows greater activity in motor processing tasks as compared to 7am, which belongs to the CON (Baker *et al*. 2018b).

In sum, all the eight cortical brain regions that we uncovered herein are clearly involved in a great diversity of cognitive tasks and are considered to be somewhat mysterious, multimodal, or integrative areas. Our findings suggest that these areas have indirect neuromodulatory properties, which may explain why they have such extensive functional profiles.

### The role of top-down arousal regulation in general fluid intelligence

Humans can reason without reliance on previously acquired knowledge. This remarkable ability is termed “fluid2 reasoning – a high-level cognitive process that extracts relational information from the environment and manipulates that information to arrive at a solution to a problem at hand. To date, no consensus theory has been established on how the brain implements this ability, which obstructs the development of techniques to improve it (e.g., via education) or prevent its decline (e.g., by eliminating harmful practices).

Fluid reasoning is often assessed using a variety of cognitive tasks – for example, sorting of abstract sensory stimuli according to a given dimension, such as shape or color (Grant and Berg 1948; Eling, Derckx and Maes 2008), or completion of a series of geometric patterns (Raven 1938), as well as tests of working and episodic memory. Processing speed and inhibitory control tasks are also used to simulate the demands of a dynamic external environment where prior knowledge cannot be utilized. The performance metrics in such seemingly disparate sets of tasks are correlated (Spearman 1904), which suggests a shared underlying neural mechanism.

Neuroscience research has identified multiple frontoparietal brain regions and their interaction playing a role in fluid reasoning (Christoff *et al*. 2001; Kroger *et al*. 2002), with mostly multiple demand, “task-positive2 network (Finn *et al*. 2015; Woolgar *et al*. 2018) and lateral prefrontal cortex in particular (Cole *et al*. 2012; Cole, Ito and Braver 2015) involved. It is not known what processes could drive such widespread network-level phenomena, but the involvement of noradrenergic modulation has been recently proposed (Tsukahara and Engle 2021). Indeed, arousal states and neuromodulatory systems are known to drive large-scale brain activity and network reconfiguration (Shine *et al*. 2019; Martin, He and Chang 2021; Oyarzabal *et al*. 2022). The results presented herein expand these ideas as follows – the properties of the ascending neuromodulatory system alone do not appear to be sufficient to explain individual variability in cognitive performance, but the properties of the whole circuit, including the top-down control of the arousal state by the PCC, do predict cognition (Figure 4).

We found that the cognition scores of older participants are less readily predicted by the PCC-arousal-RSN circuit functionality (Figure 4G). Since our dataset includes young adults only, it is difficult to bridge between these results and the published studies on arousal changes in older age. For example, it is known that cognitive abilities generally decrease with age-related changes in LC function (Bugg *et al*. 2006; Hämmerer *et al*. 2018; Dahl *et al*. 2019; Liu *et al*. 2020), however, we only observe an insignificant trend in our data that is consistent with such prior evidence (Figure 4E). This result underlies the importance of future studies considering age-related LC function as an important factor.

### Limitations

In the present study the regions driving the arousal state were calculated at the group level. However, it is possible that individual brains develop distinct connectivity pathway to control the arousal system, which could not be detectable at the group level. More detailed anatomical studies are required to discern projections from cortex to LC in humans, and a future study would be required to test functional connectivity and behavioral relevance of such arousal regulating circuits in individual brains. Nonetheless, we employed statistics (subject as a random effect) that provided evidence that the reported results generalized across the individuals in this study, and that the results are likely to generalize to other subjects not included in this study.

In the present work, the arousal state has been estimated via pupil diameter, which, in addition to LC, is driven by other neuromodulatory systems, such as the cholinergic basal forebrain (BF-ACh) (Shine *et al*. 2019). The BF-Ach system possesses general properties similar to LC-NE in the context of the present study hypothesis – that is, BF-ACh neurons branch extensively to cover the entirety of cortex (van den Brink, Pfeffer and Donner 2019), and the release of ACh in cortex also results in inhibition of spontaneous brain activity (Randić, Siminoff and Straughan 1964; Shulz *et al*. 1997). Little is known about cortical projections back to BF in humans, but animal research indicates an influence of projections from prefrontal cortex (Zaborszky *et al*. 1997). Our analysis presented in Supplementary Figure 3 reveals a complex relationship between neuomodulators and arousal, with the density of multiple receptor types predicting arousal-linked inhibition, with the strongest effect of ADRA1B noradrenergic receptor (as expected), and an opposing effect (less inhibition with higher density) by dopamine D4 density. These results confirm the complex neuromodulatory origin of pupil-linked arousal. Crucially, future research is needed to determine whether cortical regions discovered herein as arousal-driver regions influence brain state via neuromodulatory systems beyond LC.

### Implications and future directions

The present study tackles an inherently difficult problem of disentangling intrinsic mechanisms (manifesting without external intervention) that nevertheless predict cognitive performance. Why would functionality of brain circuits during resting state be predictive of task behavior? While task-related behavior is usually studied in the context of experimentally imposed variables, such behavior inevitably depends on intrinsic components as well. A crucial observation is that even the “rest2 recordings we utilize here constitute a combination of external and intrinsic factors. The experiment participants lay in a narrow tube, having their head partially fixed, keeping their eyes open, and fixating on one central point (i.e., a simplified attention task), all while making sure to remain still. To perform such “resting2, the participant must maintain a moderate level of arousal – to not fall asleep on the one hand, and not get hyperactive on the other. In addition, the participants may let their mind wander, engaging intrinsic cognitive processes such as imagery, memory, or planning. It is a possibility that the physiological mechanism for regulating a powerful neuromodulatory system that we report herein corresponds with intrinsic cognitive processes involving arousal (e.g., imagining an exciting event). While further tests are required to confirm such a hypothesis, this line of research may lead to improved understanding of generative active cognition in the wild, which is largely driven by intrinsic factors.

Another research trajectory that can be enriched by our findings is the study of how to improve human cognition and fluid reasoning in health and disease. For example, meditation is a form of mental training often requiring practicing sustained focus on an object (such as breath) which leads to improvement of cognition and, intriguingly, such training specifically involves the brain regions discovered herein as arousal-regulating regions (Tang, Hölzel and Posner 2015). Multiple studies and their meta-analysis identified a meditation-supporting network of regions that includes superior parietal cortex (region 7a), right insula and DMN regions, PCC and IPL (Tomasino *et al*. 2013). PCC activity is lower in experienced meditators during practice compared to matched controls and compared to a non-meditation-related active task (Brewer *et al*. 2011; Garrison *et al*. 2015). Mindfulness mediation also causes structural changes in the brain, for example, gray matter thickness in left PCC is smaller in experienced meditators (Berkovich-Ohana *et al*. 2020). Meta-analysis of meditation-induced connectivity shows increase of functional connectivity between PCC and AmCC (Rahrig *et al*. 2022), another region that supports arousal control according to our data. These results are generally consistent with our findings of positive correlations between these regions’ activity and arousal (Figure 3A) and the reports of stress reduction and maintenance of relaxed focus in experienced meditators. A future study is needed to determine whether meditation causally affects the large-scale arousal-related circuitry we have identified herein. Considering our finding of better cognitive performance with more negative ACME (Figure 4), a plausible hypothesis would be that meditation leads to reduction in ACME during resting state. Other options for targeting the same circuit would involve causal manipulation of the functional connections themselves, such as via invasive (Sun and Morrell 2014) or non-invasive (Sehm *et al*. 2012) brain stimulation or via neurofeedback-based training (Harmelech *et al*. 2013; Ramot *et al*. 2016, 2017).

Another important direction is to investigate the brain circuits we identified herein in neuropsychiatric disorders that involve abnormalities of the arousal system – such as Alzheimer’s, Schizophrenia, mood disorders, ADHD, and PTSD (Friedman, Adler and Davis 1999; Ressler and Nemeroff 1999; Yamamoto, Shinba and Yoshii 2014). While the typical therapeutic intervention so far entails NE regulation of brain state (Biederman and Spencer 1999; Southwick *et al*. 1999; Morilak *et al*. 2005; Nutt 2006; Campo *et al*. 2011; Moret and Briley 2011), the select higher-order cortical regions that we identified herein are likely also able to invoke modulation of brain state, and should be now considered as promising upstream regions for therapeutic interventions.

## Acknowledgements

The authors acknowledge support by the US National Institutes of Health under award R01 MH109520, and support by the US National Science Foundation under awards 2219323 and 2117429. The authors acknowledge the Office of Advanced Research Computing (OARC) at Rutgers, The State University of New Jersey, for providing access to the Amarel cluster and associated research computing resources that have contributed to the reported results. We thank Takuya Ito and members of the Cole Neurocognition Laboratory for useful comments on initial research findings.

## Supplementary figures

**Supplementary figure 1.**
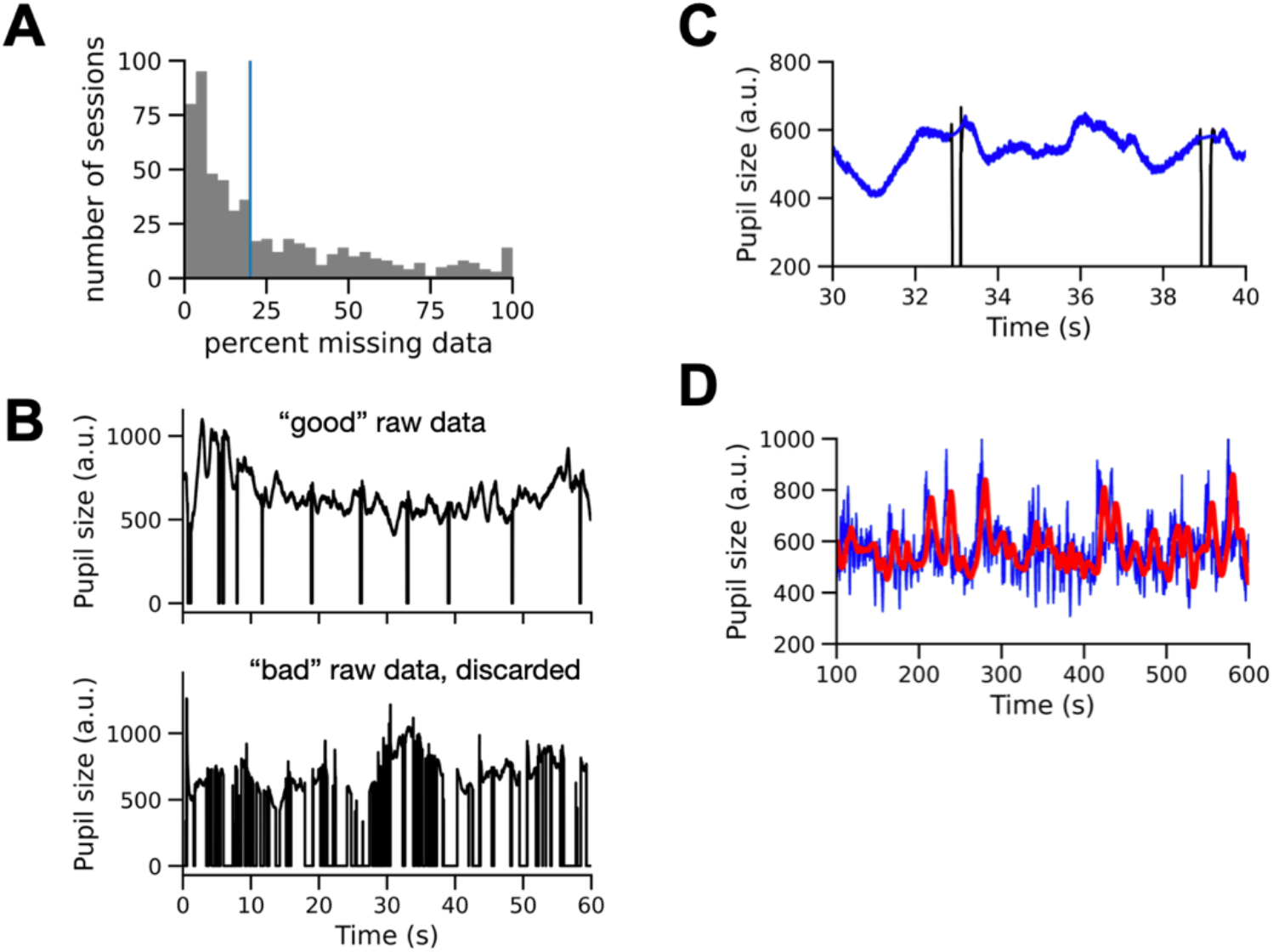
Eye-tracking data summary. **A**. Percent of time samples where the eye-tracking data were missing in each session. The blue vertical line indicates the cut-off of sessions that show above 20% missing data. **B**. Example of acceptable and unacceptable eye-tracking data quality. **C**. Blink correction procedure. Black line depicts raw pupil data. Note the drops in pupil size due to eye blinks. Blue line shows the effect of the procedure detecting the blink onset and offset and interpolating the missing points (see Methods). **D**. Example of hemodynamic response function (HRF) convolved pupil size time course. The blue line indicates pupil size time course with interpolated blinks (same as C for a more extended time period). The red line shows the result of a convolution between the pupil size time course and HRF.

**Supplementary figure 2.**
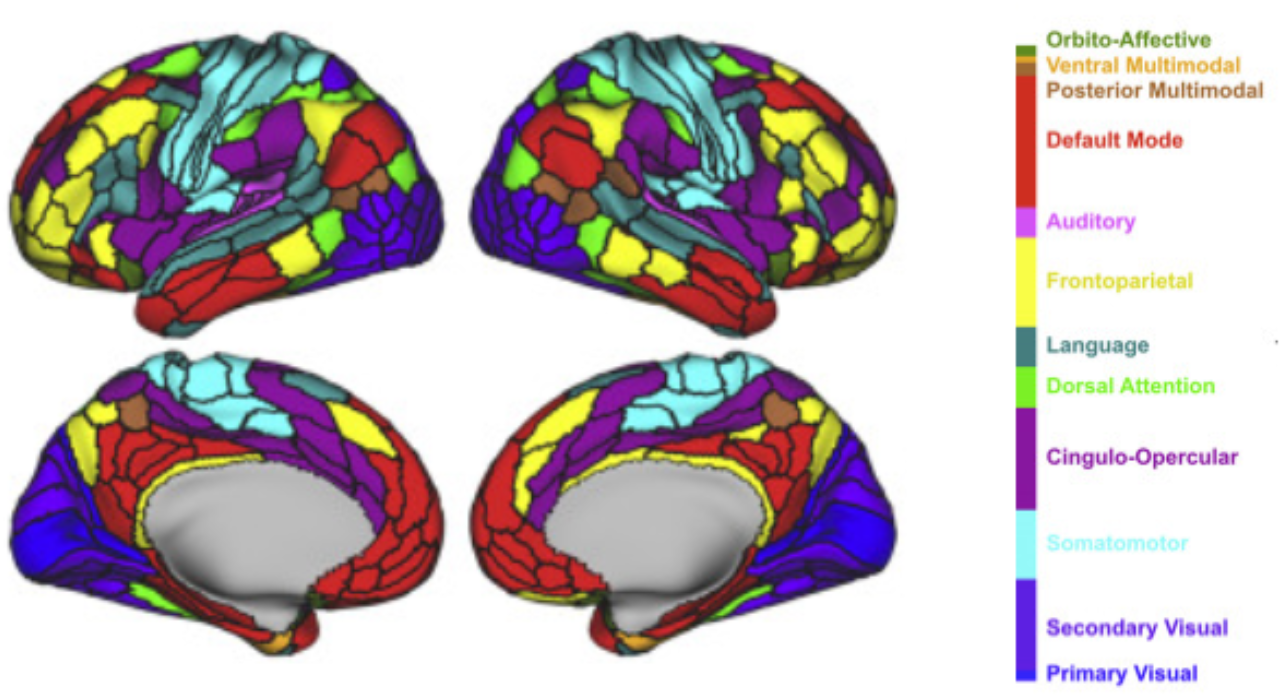
Cole-Anticevic partition. Figure reproduced with permission.

**Supplementary figure 3.**
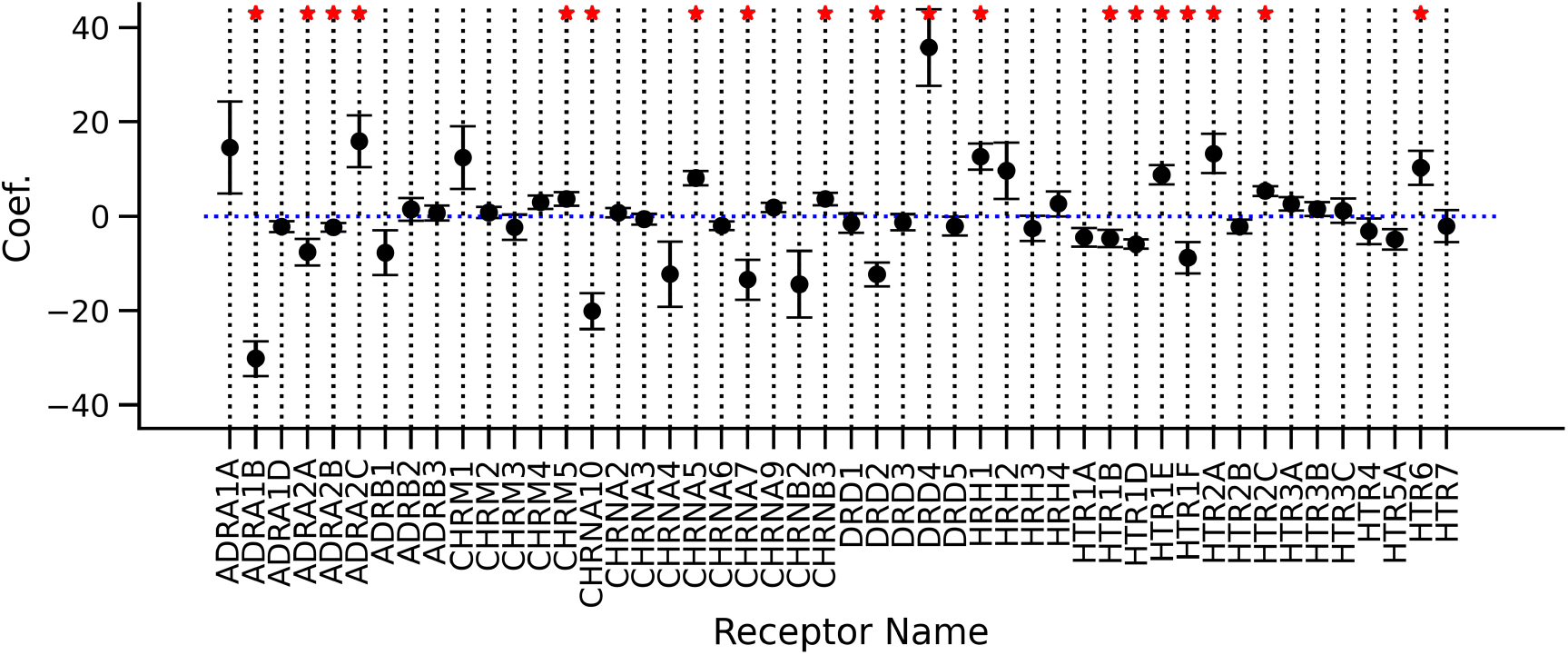
Relationship between receptor density and the arousal-linked inhibition of spontaneous brain activity. Presented are linear model coefficients indicating the power of each receptor density map at predicting arousal-linked inhibition of spontaneous brain activity on a group level. The errorbars indicate standard error of the mean and the red asterisks indicate p-value < 0.05 after FDR correction. Significant negative coefficient indicates that greater receptor density in a region predicts stronger arousal-linked inhibition of spontaneous activity, whereas a positive coefficient indicates less arousal-linked inhibition. The strong negative effect of receptor ADRA1B is most consistent with our hypothesis, with other effects indicating the complexity of neurotransmitter effects on arousal-linked changes in spontaneous brain activity. Abbreviations: ADR – Adrenalin (Norepinephrine); CHRM – Acetylcholine Muscarinic; CHRN – Acetylcholine Nicotinic; DR – Dopamine; HRH – Histamine; HTR – Serotonin;

